# Neocortical activity tracks syllable and phrasal structure of self-produced speech during reading aloud

**DOI:** 10.1101/744151

**Authors:** Mathieu Bourguignon, Nicola Molinaro, Mikel Lizarazu, Samu Taulu, Veikko Jousmäki, Marie Lallier, Manuel Carreiras, Xavier De Tiège

## Abstract

To gain novel insights into how the human brain processes self-produced auditory information during reading aloud, we investigated the coupling between neuromagnetic activity and the temporal envelope of the heard speech sounds (i.e., speech brain tracking) in a group of adults who 1) read a text aloud, 2) listened to a recording of their own speech (i.e., playback), and 3) listened to another speech recording. Coherence analyses revealed that, during reading aloud, the reader’s brain tracked the slow temporal fluctuations of the speech output. Specifically, auditory cortices tracked phrasal structure (<1 Hz) but to a lesser extent than during the two speech listening conditions. Also, the tracking of syllable structure (4–8 Hz) occurred at parietal opercula during reading aloud and at auditory cortices during listening. Directionality analyses based on renormalized partial directed coherence revealed that speech brain tracking at <1 Hz and 4–8 Hz is dominated by speech-to-brain directional coupling during both reading aloud and listening, meaning that speech brain tracking mainly entails auditory feedback processing. Nevertheless, brain-to-speech directional coupling at 4– 8 Hz was enhanced during reading aloud compared with listening, likely reflecting speech monitoring before production. Altogether, these data bring novel insights into how auditory verbal information is tracked by the human brain during perception and self-generation of connected speech.

**Highlights:**

- The brain tracks phrasal and syllabic rhythmicity of self-produced (read) speech.
- Tracking of phrasal structures is attenuated during reading compared with listening.
- Speech rhythmicity mainly drives brain activity during reading and listening.
- Brain activity drives syllabic rhythmicity more during reading than listening.

## 1. Introduction

To produce understandable speech, humans rely on self-monitoring of speech output. Such monitoring is based on neural integration of self-generated sensory information, which links speech production to speech perception (for a review, see Hickok, 2012). Still, how this self-produced sensory information is used to control speech remains unclear.

Current theories of language production consider a feedback monitoring system that monitors speech output to correct errors during production (for reviews, see Hickok, 2012; Houde and Chang, 2015). Evidence about the importance of such a system comes from adaptations of the speaker’s speech output to compensate for sensory (i.e., auditory and somatosensory) feedback manipulations (Bauer et al., 2006; Burnett et al., 1998; Guo et al., 2017; Houde, 1998; Liu et al., 2018; Shiller et al., 2009; Tremblay et al., 2003). But such feedback monitoring system cannot account for extremely fast self-corrections of speech observed in humans (Blackmer and Mitton, 1991; Nozari et al., 2011), as they require extended neural processing time. Hence, most of the current models of language production additionally include an internal speech monitoring system, which monitors speech before production. Consensus about the neural bases of such an internal system is however lacking (Gauvin et al., 2016). Indeed, some authors consider that internal speech is monitored via sensory networks similar to those involved in monitoring feedback speech (Hickok, 2012; Indefrey, 2011), while others consider that it recruits distinct neural structures such as, e.g., brain structures involved in conflict monitoring (Hickok, 2012; Nozari et al., 2011).

A potential way to gain insights into the neuronal bases of internal and feedback speech monitoring systems is to study the coupling between the speaker’s voice and its own brain activity during connected speech production. Previous magnetoencephalography (MEG) studies focusing on connected speech listening demonstrated speech-sensitive coupling between the slow modulations of the speaker’s voice and listeners’ (mainly auditory) cortex activity (Bourguignon et al., 2013; Clumeck et al., 2014; Ding et al., 2016; Gross et al., 2013; Molinaro et al., 2016; Peelle et al., 2013; Vander Ghinst et al., 2016). This coupling henceforth referred to as *speech brain tracking*, mainly occurs at syllable (4–8 Hz) and phrasal/sentential (<1 Hz) rates. It is considered to play a pivotal role in parsing connected speech into smaller units (i.e., syllables or phrases/sentences) to promote subsequent speech recognition (Park et al., 2018; Zion Golumbic et al., 2012). Additionally, it might help predict the precise timing of events in the speech stream such as syllables and phrases/sentences (Zion Golumbic et al., 2012). Such predictions probably facilitate speech comprehension as well as coordination of turn-taking transitions during verbal conversation (Friston and Frith, 2015; Zion Golumbic et al., 2012). It is then sensible to hypothesize that similar speech brain tracking is also at work during connected speech production and contribute to self-produced speech monitoring systems. If confirmed, this could bring unprecedented insights into how humans handle self-generated auditory information during language production. Additionally, investigating coupling directionality (i.e., speech → brain vs. brain → speech coupling) during connected speech production could bring critical information about the neural bases of speech production monitoring systems in humans: feedback (speech → brain coupling) vs. internal (brain → speech coupling).

To address these issues, the present MEG study relied on the comparison of speech brain tracking while subjects listened to recordings of texts read aloud (by a reader or themselves) and while they read themselves a text aloud. This approach was first chosen because previous studies from our group that investigated speech brain tracking during listening relied on live (Bourguignon et al., 2013; Clumeck et al., 2014) or recorded (Clumeck et al., 2014; Destoky et al., 2019; Vander Ghinst et al., 2019, 2016) voices continuously reading a text aloud. Second, it was also based on the shared neurocognitive processes between natural speech production and reading aloud (Sulpizio and Kinoshita, 2016). Indeed, reading aloud is recognized as a type of speech production such as, e.g., spontaneous narrative, narrative recalls, conversation, picture description (see, e.g., Bóna, 2014). The last stages of language production in those different speech situations are similar: all include phonological encoding (i.e., assigning a segment to a position in a metrical frame), phonetic encoding (i.e., retrieving the motor plans required for articulation), and articulation (i.e., producing the gestures leading to an acoustic sound) (Kawamoto et al., 2015). Settling on reading aloud also makes it possible to control speech content and linguistic form, which are two speech features previously reported to affect brain rhythms (Alexandrou et al., 2017). Reading aloud decreases the subjects’ need to focus on semantic/lexical access, other cognitive processes or speech style, which can potentially bias speech brain tracking and directionality assessments during language production (Bóna, 2014). Finally, comparing the neural processes at play during listening to somebody reading aloud and during reading aloud allows relying on auditory verbal information that shares common rhythmicity and prosody.

In practice, this MEG study investigates, using coherence and directionality analyses, speech brain tracking in subjects who (i) read a text aloud, (ii) listened to a recording of a different text, and (iii) listened to a recording of their own speech while reading aloud (i.e., playback). It was specifically designed to (i) identify cortical areas that track the slow fluctuations of self-produced speech, (ii) determine the causal nature of this tracking, and (iii) assess tracking differences between reading aloud and listening.

## 2. Methods

### 2.1. Participants

Eighteen healthy native Spanish speakers without any history of neuropsychiatric disease or language disorders were studied. One participant was excluded from the study due to excessive artifacts in the data. The study therefore reports on 17 participants (range 20–32 years; mean age 23.9 years; 9 females and 8 males). Sixteen participants were right-handed according to Edinburgh handedness inventory (score range 40–100 %; mean ± SD, 70.6 ± 19.1 %) (Oldfield, 1971). Handedness appraisal was missing from the last participant. Thirteen participants had a university degree, 1 was a master student, and 3 were trained professional with high school or secondary school degree (degree obtained at age ∼18 or ∼16 respectively when no grade is repeated). The study was approved by the BCBL Ethics Committee. Participants were included in the study after written informed consent.

### 2.2. Experimental paradigm

The experimental stimuli were derived from 2 narrative texts of ∼1000 words. The topics of the texts were maximally neutral: the first elaborated on the origin of life and human spirituality, while the second was an attempt to define what is a “discourse”. Both texts were read aloud by a male and a female native Spanish speaker and recorded with a high quality microphone. Reading pace was of 152 ± 35 words/min (mean ± SD across the four recordings).

Participants underwent four experimental conditions (*read*, *listen*, *playback*, and *rest*) lasting ∼5 minutes each while they were sitting in the MEG chair with their head inside the MEG helmet. During the *read* condition, participants continuously read aloud one of the two texts printed on A4 pages. During the *listen* condition, they listened to the audio recording of the other text read by the reader of their gender. Texts were assigned to conditions in a counterbalanced manner. During the *playback* condition, participants listened to their own voice recorded (see 2.3. for recording data acquisition details) earlier during the *read* condition. Obviously, *playback* condition was performed in all subjects after the *read* condition. This *playback* condition was used (i) to assess the impact of possible sensory prediction about upcoming speech (as subjects had some hints about speech content and production from the prior *read* condition) on speech brain tracking and on tracking directionality, and (ii) to control for potential differences in speech rhythm between *listen* and *read*. For both *listen* and *playback* conditions, sounds were played with VLC running on a MacBook pro and delivered at 60 dB (measured at ear-level in every participant) through a front-facing flat-panel loudspeaker (Panphonics Oy, Espoo, Finland) placed ∼2 m from the participants. During *rest* condition, participants were asked to fixate the gaze at a point on the wall of the magnetically shielded room (MSR) and try to reduce blinks and saccades to the minimum. The order of the conditions was either *read–listen–rest–playback* or *listen–read– rest–playback*.

### 2.3. Data acquisition

Neuromagnetic signals were recorded at the Basque Centre on Cognition, Brain and Language (BCBL) with a whole-scalp-covering neuromagnetometer installed in a MSR (Vectorview & Maxshield^TM^; MEGIN Elekta Oy, Helsinki, Finland). The 306-channel MEG sensor layout consisted in 102 sensor triplets, each comprising one magnetometer and two orthogonal planar gradiometers characterized by different patterns of spatial sensitivity to right beneath or nearby cortical sources. The recording pass-band was 0.1–330 Hz and the signals were sampled at 1 kHz. The head position inside the MEG helmet was continuously monitored by feeding current to five head-tracking coils located on the scalp and observing the corresponding coil-induced magnetic field patterns by the MEG sensors. Head position indicator coils, three anatomical fiducials, and at least 150 head-surface points (covering the whole scalp and the nose surface) were localized in a common coordinate system using an electromagnetic tracker (Fastrak, Polhemus, Colchester, VT, USA).

An optical fiber microphone was placed inside the MSR to record participants’ voice during the *read* condition. To maximize sound quality, the microphone was taped to the edge of the MEG helmet, ∼5 cm away from subjects’ mouth. Sound signals were recorded with *Audacity* at a sampling rate of 44.1 kHz. Electrooculograms (EOG) monitored vertical and horizontal eye movements, and electrocardiogram (ECG) recorded heartbeat signals. All these signals were recorded time-locked to MEG signals.

High-resolution 3D-T1 cerebral magnetic resonance images (MRI) were acquired on a 3 Tesla MRI scan (Siemens Medical System, Erlangen, Germany).

### 2.4. Data preprocessing

As reading aloud is typically associated with many sources of high-amplitude artifacts in electrophysiological signals (e.g., head movements, muscle artifacts, eye movements, etc.), special care was taken during data preprocessing to subtract as much as possible these artifacts from raw MEG data.

Continuous MEG data were first preprocessed off-line using the temporal signal space separation (tSSS) method (correlation coefficient: 0.9 and the segment length of the temporal projection set equal to the file length) to subtract external interferences, to correct for head movements, and to dampen movement artifacts induced by reading aloud (Taulu et al., 2005; Taulu and Simola, 2006). To further suppress heartbeat, eye-blink, and eye-movement artifacts, 30 independent components were evaluated from the MEG data low-pass filtered at 25 Hz using FastICA algorithm (dimension reduction, 30; non-linearity, tanh) (Hyvärinen et al., 2001; Vigario et al., 2000). Independent components displaying a correlation exceeding 0.15 with any EOG or ECG signals were subtracted from MEG data. The mean ± SD of rejected components was 7.2 ± 1.4 (*read*), 5.1 ± 1.8 (*listen*), 4.9 ± 2.0 (*rest*), and 5 ± 2.0 (*playback*). Finally, when the maximum MEG amplitude exceeded 5 pT (magnetometers) or 1 pT/cm (gradiometers), data within one second before and after the excessive amplitude were marked as artifact-contaminated to avoid analysis of MEG data compromised by any other artifact source that would not have been removed by the temporal signal space separation or independent component analysis.

Speech temporal envelopes were obtained from all sound recordings as the rectified sound signals low-pass filtered at 50 Hz. Speech temporal envelopes were further resampled at 1000 Hz time-locked to MEG signals.

### 2.5. Coherence analysis

To perform frequency and coherence analyses, continuous data obtained in all conditions (*listen*, *playback*, *read and rest*) were split into 2-s epochs with 1.6-s epoch overlap, leading to a frequency resolution of 0.5 Hz (Bortel and Sovka, 2014). MEG epochs containing periods marked as artifact contaminated were discarded from further analyses. Also, for each participant, only the minimum amount of epochs across all conditions was used for subsequent analyses. These steps led to 703±45 artifact-free epochs of MEG and voice envelope signals for each participant and condition.

Coherence is an extension of Pearson correlation coefficient to the frequency domain that determines the degree of coupling between two signals, providing a number between 0 (no linear dependency) and 1 (perfect linear dependency) for each frequency (Halliday, 1995). Coherence was previously used to assess the coupling between voice and brain signals at the frequencies corresponding to phrasal/sentential (<1 Hz) and syllable (4–8 Hz) rates (Bourguignon et al., 2013; Luo and Poeppel, 2007; Molinaro and Lizarazu, 2017; Peelle et al., 2013; Poeppel, 2003; Vander Ghinst et al., 2016).

Coherence was first estimated at the sensor level. Data from gradiometer pairs were combined in the direction of maximum coherence as done in Bourguignon et al. (2015). Coherence at phrasal/sentential level was taken at the frequency bin corresponding to 0.5 Hz, and coherence at syllable level was taken as the mean across coherence at frequency bins comprised in 4–8 Hz.

Coherence was also evaluated at the source level using a beamformer approach since this method has a high sensitivity to activity coming from locations of interest while attenuating external interferences such as reading-induced head movement, eye movements, or muscle artifacts (Hillebrand et al., 2005). To do so, individual MRIs were first segmented using Freesurfer software (Martinos Center for Biomedical Imaging, Massachusetts, USA; Reuter et al., 2012). Then, the MEG forward model was computed for three orthogonal tangential current dipoles placed on a homogeneous 5-mm grid source space that covered the entire brain (MNE suite; Martinos Center for Biomedical Imaging, Massachusetts, USA; Gramfort et al., 2014) and further reduced to its two first principal components. Finally, coherence maps were produced within the computed source space at 0.5 Hz and 4–8 Hz using Dynamic Imaging of Coherent Sources (DICS) (Gross et al., 2001), and further interpolated onto a 1-mm grid. Both planar gradiometers and magnetometers were used for inverse modeling after dividing each sensor signal by its noise variance. Despite the fact that raw magnetometer signals are considered noisier than planar gradiometers, in the framework of signal space separation, signals from both sensor types are reconstructed from the same inner components, corresponding to the magnetostatic multipole expansion, and have therefore similar levels of residual interference after suppression of signals from external sources (Garcés et al., 2017). This explains why both sensor types were used for source reconstruction. The noise variance was estimated from the continuous *rest* MEG data band-passed through 1–195 Hz, for each sensor separately. As the analyses described in a further paragraph require extracting the time course of some sources, we used the additional constraint that beamformer weight coefficients are real-valued. This constraint is sensible since one can easily argue that electrical currents in the brain are real--valued. Practically, it leads to using the real part of the cross-spectral density matrix in DICS beamformer computation.

To compute group-level coherence maps, a non-linear transformation from individual MRIs to the standard Montreal Neurological Institute (MNI) brain was first computed using the spatial-normalization algorithm implemented in Statistical Parametric Mapping (SPM8, Wellcome Department of Cognitive Neurology, London, UK; Ashburner et al., 1997; Ashburner and Friston, 1999) and then applied to individual MRIs and coherence maps. This procedure generated a normalized coherence map in the MNI space with 1-mm cubic voxels for each subject, condition and frequency of interest (i.e., 0.5 Hz and 4–8 Hz). Group-level maps were obtained by averaging the normalized coherence maps across participants and conditions.

### 2.6. Directionality assessment

The directionality of the coupling between the voice signals and the activity within brain areas displaying a significant local maximum of coherence (see 2.8.), was assessed with renormalized partial directed coherence (rPDC) (Schelter et al., 2009, 2006). To this aim, the time-course of brain electrical activity within these brain areas was estimated with the beamformer described in 2.5., in the direction maximizing the coherence with speech temporal envelope. Source and voice signals were low-pass filtered at 10 Hz and down-sampled at 20 Hz. Then, for each source separately, a vector autoregressive (VAR) model of order 40 was fitted to the source and the voice data using the ARfit package (Schneider and Neumaier, 2001). The rPDC was then estimated based on the Fourier transform of the VAR model coefficients. This enabled for estimating rPDC at frequencies from 0 to 10 Hz with 0.5 Hz resolution.

### 2.7. Partial coherence to control for artifacts

In the *read* condition, there was a discrepancy between sensor and source-level results (see Results section). In the sensor space, strong artifacts at the edge of the sensor array obscured the 4–8-Hz speech brain tracking. In the source space, artifacts were present but genuine speech brain tracking in auditory cortices was clearly visible thanks to the use of the beamformer approach. To verify that this discrepancy pertained to that beamformer did effectively dampen artifacts—and hence strengthen results derived from source-space data—, we estimated the coherence between speech temporal envelope and MEG signals while partialling out the contribution of MEG signals recorded at sensors on the edge of the sensor array.

The following analysis was performed separately at 0.5 Hz and 4–8 Hz. For each gradiometer pair on the edge of the sensor array (23 in total), we estimated the orientation in the 2-d space spanned by both gradiometer signals (Bourguignon et al., 2015) yielding the maximum coherence with speech temporal envelope. Partial coherence was then estimated between speech temporal envelope and all gradiometer signals (again optimizing on the orientation within all pairs) while partialling out edge gradiometer signal in its optimal orientation (Halliday, 1995). This led to as many sensor distribution of partial coherence as there are edge gradiometer pairs. For each sensor, we retained the minimum partial coherence value across all these edge gradiometer pairs.

### 2.8. Statistical analyses

#### 2.8.1 Reading pace

The word per minute rate produced in the *read* condition by the participants was compared to the one of the texts used in the *listen* condition with a paired *t*-test.

#### 2.8.2. Significance of subject-level coherence in the sensor space

We evaluated the statistical significance of sensor-space coherence values, using surrogate-data-based statistics (Faes et al., 2004). For each participant, condition, and frequency range of interest (i.e., 0.5 Hz and 4–8 Hz), we extracted the maximum across gradiometer pairs of the mean coherence across the frequency range of interest. This maximum genuine coherence was then compared to a distribution of 1000 surrogate values computed in the same way, but with speech temporal envelope replaced by its Fourier transform surrogate (Faes et al., 2004). Fourier transform surrogate preserves the power spectrum but destroys the phase information by replacing the phase of Fourier coefficients by random numbers in the range [–π; π] (Faes et al., 2004). Genuine maximum coherence values were deemed significant when they exceeded the 95^th^ percentile of their surrogate distribution.

#### 2.8.3. Significance of group-level coherence in the source space

The statistical significance of group-level coherence maps was assessed with non-parametric permutation test. First, participant- and group-level *rest* coherence maps at the frequencies of interest (i.e., 0.5 Hs and 4–8 Hz) were computed with *rest* MEG and voice (of *read* and *listen* conditions) signals. Group-level difference maps were obtained by subtracting *f*-transformed genuine (*read*, *listen* or *playback*) and *rest* group-level coherence maps for each frequency of interest. Under the null hypothesis that coherence maps are the same whatever the experimental condition, the labeling genuine or *rest* are exchangeable prior to difference map computation (Nichols and Holmes, 2002). To reject this hypothesis and to compute a significance threshold for the correctly labeled difference map, the sample distribution of the maximum of the difference map’s absolute value within the preselected brain areas was computed from a subset of 1000 permutations. The threshold at *p* < 0.05 was computed as the 95 percentile of the sample distribution (Nichols and Holmes, 2002). All supra-threshold local coherence maxima were interpreted as indicative of brain regions showing statistically significant coupling with the produced (*read*) or heard (*listen* and *playback*) sounds.

#### 2.8.4. Comparison of source location between conditions

The coordinates of significant local coherence maxima were statistically compared between conditions (*listen* vs. *playback*, *listen* vs. *read*, and *playback* vs. *read*) using the location-comparison approach proposed by Bourguignon et al. (2017). This method uses a bootstrap procedure (Efron, 1979) to estimate the sample distribution of coordinates of the two local coherence maxima under comparison and tests the null hypothesis that the distance between them is zero. Briefly, we generated 1000 group-level maps of the conditions under assessment by random bootstrapping from the individual maps, and identified the coordinates of the local maxima closest to the genuine maxima location. The resulting sample distribution of coordinate difference was then submitted to a multivariate location test evaluating the probability that this difference is zero (Bourguignon et al., 2017). That test tightly relates to the multivariate *T*^2^ test (Hotelling, 1931) and assumes that the sample distribution of coordinates difference is normal.

For one local maximum, we further tested the—*a posteriori*—hypothesis that its bootstrap coordinate distribution was bimodal rather than unimodal, suggesting that two separate sources would contribute to that single local maximum. As a first step, we built a map of bootstrap source density with 1-mm cubic voxels, which we will denote *D*(*r*) with *r* = (*x*,*y*,*z*) indexing voxels. *D*(*r*) was initially set to be uniformly 0, and for each bootstrap source coordinate, we added a value 1 at the corresponding voxel. *D*(*r*) was further smoothed with a 5-mm FWHM gaussian kernel. We then used matlab *fminsearch* function to fit two models to *D*(*r*): a Gaussian distribution, and a mixture of 2 Gaussian distributions. Formally, the first model was

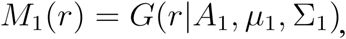

and the second model was

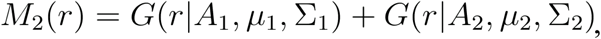

where

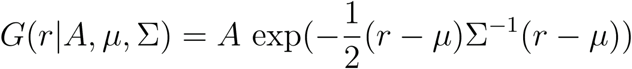

is a 3-d Gaussian distribution with *A* its amplitude, 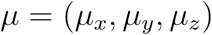 its center, and

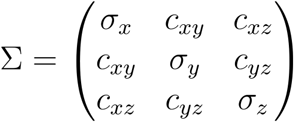

its—symmetric—covariance matrix. Hence, there were *df*_1_ = 10 parameters in *M*_1_(*r*) and *df*_2_ = 20 in *M*_2_(*r*). We then used a Fisher test to compare statistically the proportion of variance explained by these two models. These proportions can be written as 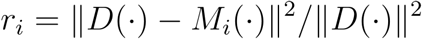, with 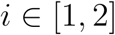 and 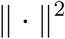 the sum of squares across all voxels. Under the null hypothesis that *M*_2_ does not do any better than *M*_1_, the quantity

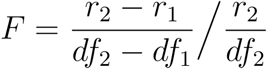

follows a *F* distribution with *df*_1_ and *df*_2_ degrees of freedom. This null hypothesis can be disproved if *F* exceeds the percentile 95^th^ of *F*_10,20_.

#### 2.8.5. Significance of individual subjects’ rPDC values and comparison between coupling directions

We evaluated the number of participants showing statistically significant rPDC, using surrogate-data-based statistics (Faes et al., 2010). Statistical analyses were performed on rPDC at 0.5 Hz or 4–8 Hz depending on whether the source was identified on phrasal/sentential- or syllable-level coherence map. For each participant, selected brain area, and coupling direction, the genuine rPDC value (at 0.5 Hz or the mean across 4–8 Hz) was compared to a distribution of 1000 surrogate rPDC values derived from causal Fourier transform surrogate data (Faes et al., 2010). Causal Fourier transform surrogate data were generated with the estimated VAR model wherein coupling in the specific causal direction being tested is abolished by setting to 0 the associated coefficients. Genuine rPDC values were deemed significant when they exceeded the 95^th^ percentile of their surrogate distribution.

Values of rPDC were compared between speech → brain and brain → speech directions using paired t-tests across participants.

#### 2.8.6. ANOVA assessment of coherence, rPDC, and partial coherence values

Source-level coherence, rPDC and sensor-level partial coherence values were analyzed with 2-way repeated measures ANOVAs. In these assessments, the factors were the condition (*listen*, *playback*, and *read*), and the sensor/source location. ANOVAs were run separately for 0.5 Hz and 4–8 Hz coupling, and for speech → brain and brain → speech directions in case of rPDC assessment. This is justified by that coupling values within these two classes had relatively different variances. Analysing data together would have violated the homoscedasticity assumption of the ANOVA. For source-level coherence values, the dependent variable was the maximum coherence across a 10-mm sphere centered on significant local maxima of group-level coherence maps. For sensor-level partial coherence values, the dependent variable was the maximum partial coherence across subsets of gradiometer pairs showing the peaks of coherence. Formally, these subsections comprised the 9 gradiometers of maximum coherence averaged across participants and conditions. There were 2 selections, one for the left and one for the right hemisphere.

### 2.9. Data and software availability

Data and analysis scripts are available upon reasonable request to the corresponding author.

## 3. Results

### 3.1 Reading pace

In the *read* condition, participants read at a pace of 158 ± 17 words per min (mean ± SD). This pace was not significantly different from the one they heard in the *listen* condition (*t*_16_ = 1.26, *p* = 0.23).

### 3.2 Coherence results

#### 3.2.1 Coherence in the sensor space

Figure 1 illustrates the results of speech brain tracking quantified with coherence in the sensor space. The maximum coherence between MEG signals and speech temporal envelope peaked at 0.5 Hz and at 4–8 Hz. These frequency ranges match the supra-second phrasal/sentential time-scale (0.5 Hz) and the 150–300-ms syllable time-scale (4–8 Hz). In both listening conditions (*listen* & *playback*), the topography of the coherence was characterized by clusters over bilateral posterior temporal sensors. In the *read* condition, coherence topographies were suggestive of the presence of strong artifacts but also of genuine bilateral activity arising from posterior temporal sensors (more convincingly so at 0.5 Hz than at 4–8 Hz).

**Figure 1.**
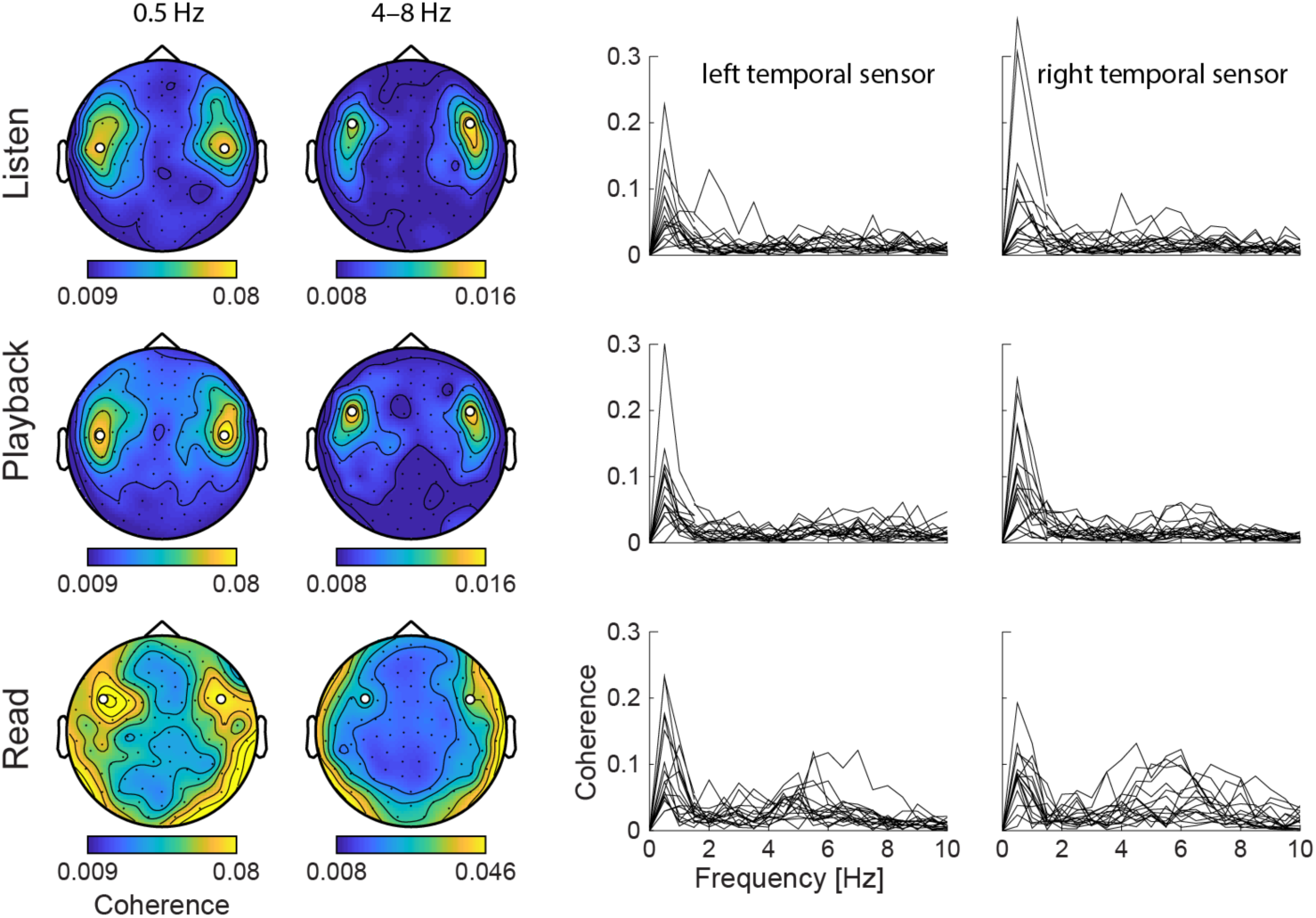
Coherence at the sensor level. *Left*—Sensor distribution of the coherence at 0.5 Hz and 4–8 Hz averaged across subjects. White discs highlight the sensors of maximum coherence, or, in the read condition at 4–8 Hz, the sensors suggestive of the presence of genuine speech brain tracking. *Right*—Individual coherence spectra at the highlighted sensors. Values from 0 to 1.5 Hz are taken from sensors identified in the 0.5 Hz map, and values from 1.5 Hz to 10 Hz from the sensors identified in the 4–8 Hz map.

Coherence in the sensor space was significant in all participants and conditions at 0.5 Hz, and in 13 (*listen*), 12 (*playback*), and 17 (*read*) out of 17 participants at 4–8 Hz. Note that the detection rate of significant coherence in the *read* condition has likely been inflated by the presence of artifacts inherent to speech production.

#### 3.2.2 Coherence in the source space

Figure 2A presents the source-space coherence maps obtained with DICS at 0.5 Hz and 4–8 Hz separately.

**Figure 2.**
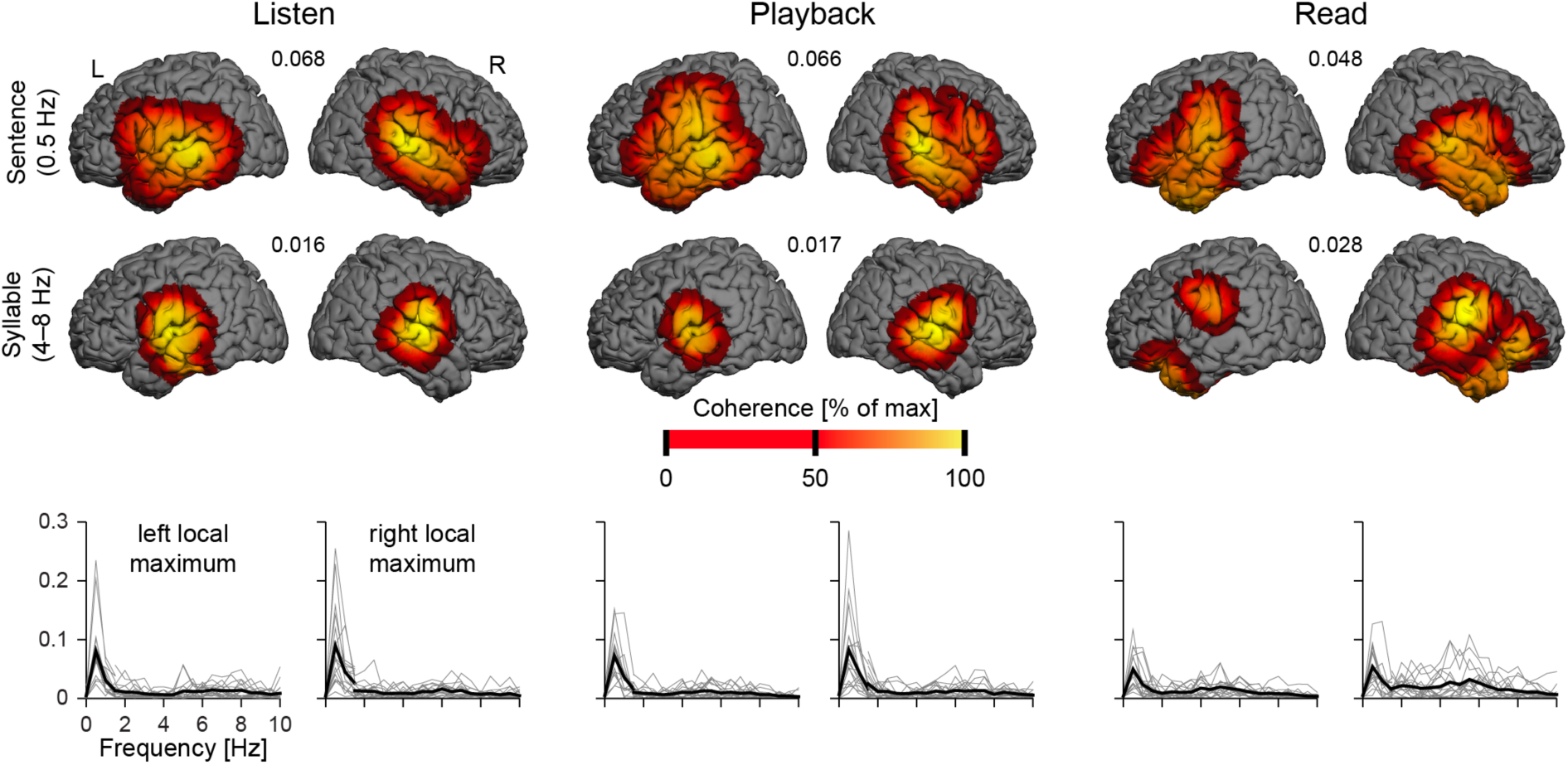
Coherence in the source space. *Top*—Group-level coherence maps at 0.5 Hz and 4– 8 Hz in the 3 conditions (*listen*, *playbach* and *read*) thresholded at statistical significance level. The color scale is tailored to each coherence map: it ranges from 0 to its maximum (indicated in between brain images). *Bottom*—Individual (gray) and group-averaged (black) coherence spectra at the local maxima of coherence.

Table 1 presents the MNI coordinates of significant local coherence maxima observed in source-space maps.

**Table 1.** MNI coordinates [mm] an coherence values of maximum speech brain tracking, as well as corresponding sensor-level coherence values controlled for artifacts in sensors at the edge of the sensor array.

|  | Left hemisphere |  |  | Right hemisphere |  |  |
| --- | --- | --- | --- | --- | --- | --- |
|  | MNI coordinate [mm] | Source coherence | Sensor partial coherence | MNI coordinate [mm] | Source coherence | Sensor partial coherence |
| Speech brain tracking at 0.5 Hz |  |  |  |  |  |  |
| *listen* | [-66 -25 1] | 0.068 | 0.056 | [66 -25 7] | 0.070 | 0.060 |
| *playback* | [-67 -28 -3] | 0.063 | 0.046 | [66 -24 3] | 0.068 | 0.046 |
| *read* | [-62 -10 12] | 0.040 | 0.045 | [66 -22 6] | 0.043 | 0.041 |
| Speech brain tracking at 4–8 Hz |  |  |  |  |  |  |
| *listen* | [-61 -12 7] | 0.0159 | 0.0138 | [65 -13 7] | 0.0162 | 0.0133 |
| *playback* | [-63 -12 9] | 0.0153 | 0.0122 | [65 -11 7] | 0.0172 | 0.0135 |
| *read* | [-62 -13 28] | 0.0209 | 0.0174 | [65 -10 18] | 0.0286 | 0.0249 |

In both listening conditions (*listen* & *playback*) significant local coherence maxima localized in bilateral cortex around posterior superior temporal sulcus (pSTS) at 0.5 Hz and in bilateral supratemporal auditory cortex (STAC) at 4–8 Hz. The location comparison test revealed no statistically significant difference in location between these two conditions (*ps* > 0.5; 4 comparisons: 2 frequencies × 2 hemispheres).

In the *read* condition, source reconstruction results emphasized the presence of genuine speech brain tracking. Some artifacts remained that peaked nearby the pons (0.5 Hz, MNI [–1 –1 –35], coherence 0.049; 4–8 Hz, MNI [2 –14 –36], coherence 0.028), but they did not overshadow coherence local maxima related to genuine speech brain tracking (see Figure 2A and Table 1 for peak coordinates and coherence values).

The speech brain tracking elicited by the *read* condition appeared to be different from that during listening conditions at both 0.5 Hz and 4–8 Hz. We focus below on the comparison between *read* and *listen*, but similar results were obtained from the comparison between *read* and *playback*.

At 0.5 Hz, right-hemisphere local coherence maxima in *read* and *listen* were distant of only 3 mm, a distance that was not statistically significant (*F*_3,998_ = 0.052, *p* = 0.98). In the left hemisphere, they were distant of 19 mm, which, surprisingly, was not deemed statistically significant either (*F*_3,998_ = 1.41, *p* = 0.24). Detailed analyses revealed that this lack of significance pertained to that coordinates of local coherence maxima in the *listen* condition had a bimodal — rather than unimodal — distribution, which hampered the location-comparison test. Indeed, maps of source density revealed that coherence in the *listen* condition peaked mainly at pSTS ([–66 –27 1]) but also at STAC ([–64 –13 6]). Also, a model with 2 Gaussian distributions explained 99.90% of the variance of the source density map, which was significantly better than the 95.76% explained by a model based on a single Gaussian distribution (*F*_10,20_ = 7.40, *p* < 0.0001). In the 2-Gaussian model, individual distributions were centered on [–66.3 –27.5 1.1] and [–64.0 –15.1 5.5]. Relative importance of the two Gaussian distributions 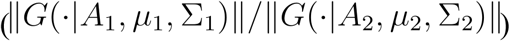 was 5.3, indicating that group-level coherence in the *listen* condition peaked ∼5.3 times more often in the first than in the second cluster. Also, the center of this second cluster was only 8.6 mm away from the maximum in the *read* condition. Of notice, there was only one peak in the source density map of the *read* condition. These results indicate that reading aloud elicits speech brain tracking only in STAC while speech listening also recruits the cortex around the pSTS.

At 4–8 Hz, local coherence maxima in the *read* condition localized in bilateral parietal operculum, i.e., more dorsally (above the sylvian fissure) than those in the *listen* condition by 19 mm (left hemisphere) and 11 mm (right hemisphere). The location-comparison test confirmed that this difference in location between *read* and *listen* conditions was statistically significant (left hemisphere, *F*_3,998_ = 10.10, *p* < 0.0001; right hemisphere, *F*_3,998_ = 3.49, *p* = 0.015).

#### 3.2.3 Effect of conditions on the coherence strength

Speech brain tracking values quantified with coherence at condition-specific dominant sources were compared with repeated measures ANOVA, separately at 0.5 Hz and 4–8 Hz.

At 0.5 Hz there was a main effect of condition on coherence level (*F*_2,32_ = 8.10, *p* = 0.0014), no significant main effect of hemisphere (*F*_1,16_ = 0.20, *p* = 0.66), and no significant interaction (*F*_2,32_ = 1.95, *p* = 0.16). Post-hoc t-tests revealed that coherence values in *listen* (0.092 ± 0.039, mean ± SD of the mean coherence across hemispheres) and *playback* (0.090 ± 0.046) did not differ significantly (*t*_16_ = 0.21, *p* = 0.84), while values in *read* (0.057 ± 0.022) were significantly lower than those in *listen* (*t*_16_ = 3.95, *p* = 0.0012) and *playback* (*t*_16_ = 3.47, *p* = 0.0031).

At 4–8 Hz there was a main effect of condition on coherence level (*F*_2,32_ = 16.6, *p* < 0.0001), no significant main effect of hemisphere (*F*_1,16_ = 2.23, *p* = 0.15), and no significant interaction (*F*_2,32_ = 0.06, *p* = 0.94). Post-hoc t-tests revealed that coherence values in *listen* (0.0183 ± 0.0052) and *playback* (0.0191 ± 0.052) did not differ significantly (*t*_16_ = 0.58, *p* = 0.57), while values in *read* (0.0294 ± 0.0086) were significantly higher than those in *listen* (*t*_16_ = 4.28, *p* = 0.0006) and *playback* (*t*_16_ = 4.37, *p* = 0.0005).

### 3.3. Directionality results

rPDC was used to separate the relative contributions to speech brain tracking of signals reacting to speech (i.e., external feedback monitoring system) and signals preceding speech (i.e., internal speech monitoring system).

Figure 3 presents rPDC values in all conditions.

**Figure 3.**
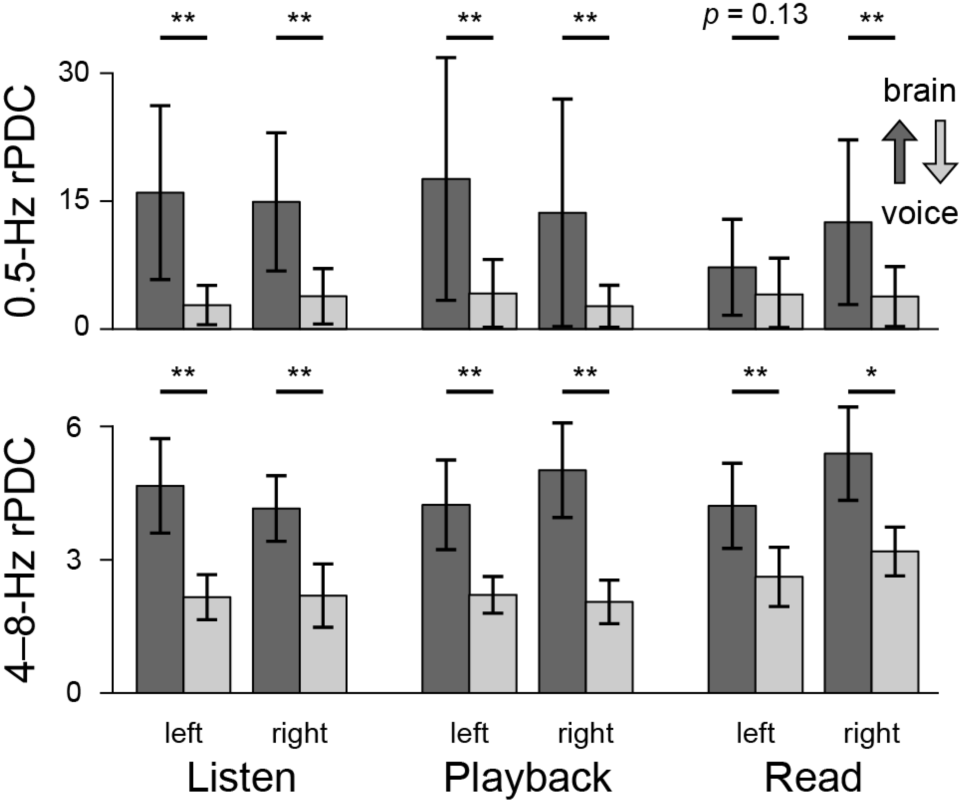
Directionality assessment with renormalized partial directed coherence (rPDC). Bars display the mean and SD of rPDC values. There is one bar per conditions (*listen*, *playback* and *read*), frequency range of interest (0.5 Hz and 4–8 Hz), hemisphere (left and right), and direction (speech → brain and brain → speech). Significance of the comparison between directions are indicated above each pair of bars (* *p* < 0.05, ** *p* < 0.01).

Table 2 details the number of participants displaying significant rPDC in all conditions, directions and frequency of interest.

**Table 2.** Number of subjects displaying significant renormalized partial directed coherence (rPDC).

|  |  | *listen* |  | *playback* |  | *read* |  |
| --- | --- | --- | --- | --- | --- | --- | --- |
|  |  | left | right | left | right | left | right |
| 0.5 Hz | speech → brain | 16 | 15 | 14 | 13 | 10 | 12 |
|  | brain → speech | 4 | 5 | 5 | 3 | 5 | 4 |
| 4–8 Hz | speech → brain | 10 | 8 | 9 | 9 | 12 | 9 |
|  | brain → speech | 0 | 0 | 1 | 1 | 4 | 6 |

Paired t-tests revealed that rPDC was systematically higher in the speech → brain direction than in the brain → speech direction (*ps* < 0.05) except at 0.5 Hz in the left hemisphere in the *read* condition (*t*_16_ = 1.61, *p* = 0.13).

The ANOVA assessment of rPDC values was performed with factors condition (*listen*, *playback* and *read*) and hemisphere (left and right) separately at 0.5 Hz and 4–8 Hz, and for the two coupling directions. There was a significant main effect of condition on speech → brain rPDC at 0.5 Hz (*F*_2,32_ = 4.66, *p* = 0.017) explained by that values in *read* (10.8 ± 7.2, mean ± SD of the mean rPDC across hemispheres) were lower than those in *listen* (16.9 ± 7.9; *t*_16_ = 2.70, *p* = 0.016) and *playback* (17.0 ± 11.9; *t*_16_ = 3.45, *p* = 0.0033), while the two latter did not differ significantly (*t*_16_ = 0.063, *p* = 0.95). There was also a significant effect of condition on brain → speech rPDC at 4–8 Hz (*F*_2,32_ = 8.43, *p* = 0.0011) explained by that values in *read* (2.75 ± 0.74) were higher than those in *listen* (2.06 ± 0.38; *t*_16_ = 2.90, *p* = 0.011) and *playback* (2.02 ± 0.38; *t*_16_ = 3.50, *p* = 0.0030), while two latter did not differ significantly (*t*_16_ = 0.30, *p* = 0.77). There were no other significant main effects or interactions (*ps* > 0.1).

As it is unclear how artifacts contributed to these results, we repeated the rPDC analysis between speech temporal envelope and signals from a sensor that picked up strong artifacts (left hemisphere: MEG153*; right hemisphere: MEG263*). The ANOVA assessment of these rPDC values revealed in all 4 instances (2 coupling directions × 2 frequency ranges) a significant effect of condition (*ps* < 0.05) explained by higher values in *read* than in *listen* and *playback*.

### 3.4. Partial coherence

Figure 4 illustrates speech brain tracking in sensor space controlled for artifacts in edge sensors using partial coherence. It is noteworthy that in *read* condition, artifacts were substantially suppressed by using partial coherence, while coherence at bilateral auditory cortices was essentially preserved. Moreover, partial coherence values were quite faithful to the source-space coherence values, as can be seen in group-level values displayed in Table 1 (similarity in source coherence and sensor partial coherence values).

**Figure 4.**
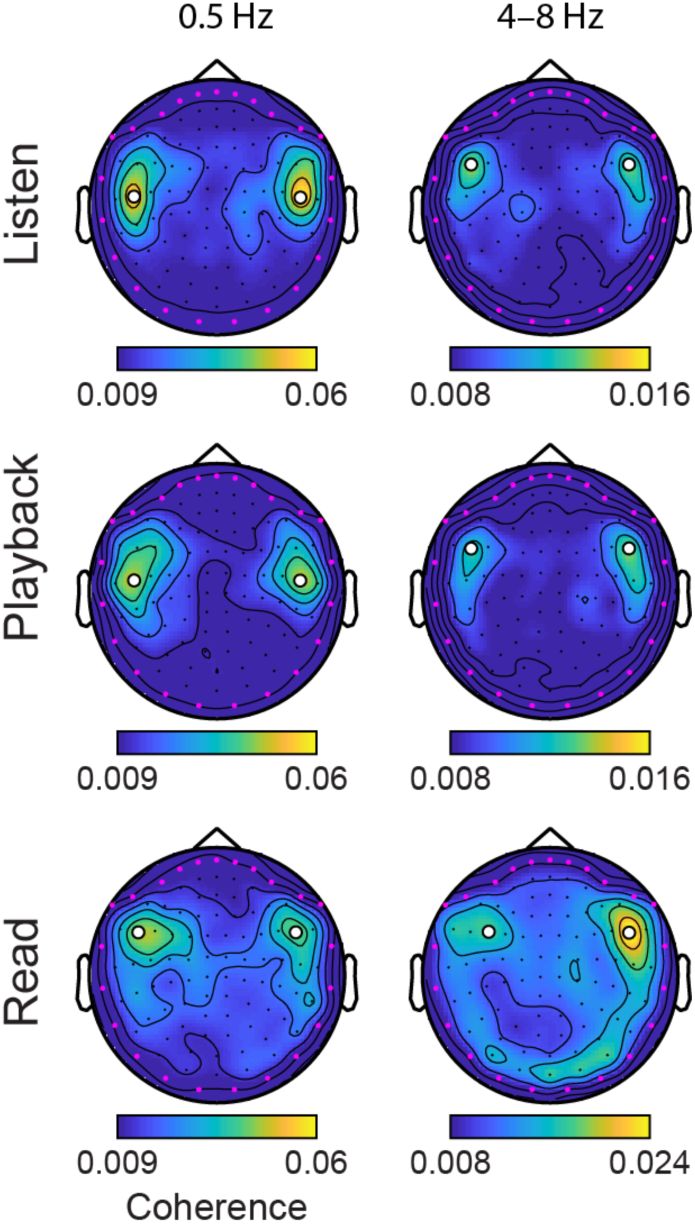
Speech brain tracking at the sensor level assessed with partial coherence to control for artifacts in edge sensors (highlighted in magenta). Note that topographies at 4–8 Hz are displayed with a different scale for *read* and listening (*listen* and *playback*) conditions. White discs highlight the same sensors as those in figure 1. sensors of maximum coherence, or, in the read condition at 4–8 Hz, the sensors suggestive of the presence of genuine speech brain tracking.

Partial coherence levels were compared with repeated measures ANOVA with factors condition (*listen*, *playback* and *read*) and hemisphere (left and right) separately at 0.5 Hz and 4–8 Hz. At 0.5 Hz, there were no significant effects nor interaction (*ps* > 0.5). At 4–8 Hz there was a main effect of condition (*F*_2,32_ = 18.3, *p* < 0.0001), no significant main effect of hemisphere (*F*_1,16_ = 1.27, *p* = 0.28), and no significant interaction (*F*_2,32_ = 0.57, *p* = 0.57). Partial coherence values in *read* (0.0292 ± 0.0106, mean ± SD of the mean coherence across hemispheres) were higher than those in *listen* (0.0157 ± 0.0049; *t*_16_ = 4.38, *p* = 0.0005) and *playback* (0.0158 ± 0.0046; *t*_16_ = 4.41, *p* = 0.0004), while two latter did not differ significantly (*t*_16_ = 0.14, *p* = 0.89).

## 4. Discussion

This study demonstrates that during reading aloud, the speaker’s brain tracks the slow temporal fluctuations of speech output. The auditory cortex tracks sentence/phrase structure (<1 Hz) while parietal operculum tracks syllable structure (4–8 Hz). It also brings novel insights into the neural bases of speech production monitoring systems while reading aloud.

### 4.1. Speech brain tracking at frequencies <1 Hz

We found that <1-Hz speech brain tracking was attenuated during self-produced speech compared with listening to external speech. A control analysis, however, failed to corroborate this finding as it indicated similar rather than lower level of <1-Hz tracking during reading compared with listening. An attenuation would be well in line with the literature. Indeed, auditory cortical responses (i.e., N100/M100 evoked response) to self-produced speech are typically attenuated or suppressed compared with those obtained during listening to a playback recording of the same sounds or during silent reading of a text (Curio et al., 2000; Houde et al., 2002; Numminen et al., 1999; Numminen and Curio, 1999). Such attenuation is absent when the auditory feedback is altered (e.g., pitch-shifted or alien speech feedback) (Heinks-Maldonado et al., 2006, 2005).

Our results also indicate that <1-Hz speech brain tracking while reading aloud is dominated by the speech feedback monitoring system. Indeed, both reading and listening gave rise to similarly low level of <1-Hz brain → speech coupling, which we posit, is the hallmark of reliance on forward models. Note that the significant brain → speech coupling observed in ∼30% of the subjects was most likely spurious, i.e., related to the fact that, in directionality assessment, strong coupling in one direction generates spurious coupling in the other direction (Faes et al., 2010).

Our results also shed light on the neural network involved in monitoring <1-Hz fluctuations in speech temporal envelope. During speech listening, this network seems to include the STAC and cortex around pSTS, while it only involves the STAC during reading aloud. This suggests that during self-generated speech, sensory feedback at phrasal/sentential level is mainly processed at early auditory cortices.

### 4.2. Speech brain tracking at 4–8 Hz

At 4–8 Hz, speech brain tracking was stronger when reading aloud than during passive listening and it peaked in different cortical areas, i.e., STAC during listening and parietal operculum during reading aloud. Tracking was mainly driven by the speech → brain contribution during reading aloud similarly to the listening conditions. There was however a significant enhancement in brain → speech coupling during reading compared with listening conditions.

In humans, speech temporal envelope essentially fluctuates at 2–10 Hz, peaking at ∼5 Hz (Ding et al., 2017). This corresponds to the mean syllable rate of speech (5–8 Hz) across many languages (Pellegrino et al., 2011). These findings led some authors to consider that this frequency range likely indicates universal rhythmic properties of human speech constrained by the neural dynamics of speech production/perception and the biomechanical properties of human articulator (Ding et al., 2017). Interestingly, a previous MEG study demonstrated the existence of significant coupling between ventral primary sensorimotor (SM1) cortex (i.e., mouth area) and orbicular oris muscle activities during silent mouthing of a syllable (/pa/) periodically repeated at different frequencies (i.e., 0.8–5 Hz) (Ruspantini et al., 2012). This coupling phenomenon was driven by the mouth movement repetition rate during syllable mouthing and peaked at the individual spontaneous movement rate (i.e., self-paced rate of syllable articulation: ∼2–3 Hz). It is therefore probably analogous (for a detailed discussion, see Bourguignon et al., n.d.) to the previously described cortico-kinematic coherence (CKC) phenomenon, which is the coupling between the kinematics of finger or toe movements and the activity in the SM1 cortex corresponding to the moved limb (Bourguignon et al., 2012, 2011; Marty et al., 2015; Marty et al., 2015; Piitulainen et al., 2015). CKC indeed occurs at movement frequency (and harmonics), which is rather similarly visible in the rectified surface electromyogram and other kinematic-related signals such as acceleration, force and pressure (Piitulainen et al., 2013). Of note, CKC is mainly driven by proprioceptive afferents to SM1 cortex (Bourguignon et al., 2015; Piitulainen et al., 2013). Accordingly, our data suggests that during connected speech production, self-generated proprioceptive and auditory information resulting from syllable production are monitored in ventral SM1 cortex. In particular, the multimodal (i.e., somatosensory and auditory) nature of such speech-related sensory monitoring at SM1 cortex is supported by the rather low correlation between rhythmical lip movement and auditory speech temporal envelope during speech production (see, e.g., Bourguignon et al., 2018; Chandrasekaran et al., 2009; Park et al., 2016). The observed frequency-specific auditory feedback monitoring at SM1 cortex is in agreement with the external feedback monitoring system and the sensorimotor transformation theories of speech (Cogan et al., 2014; Hickok, 2012; Houde and Chang, 2015). Critically, the present study suggests that the neocortical areas involved in 4–8 Hz speech brain tracking are different during speech perception and production, which brings novel major insights into the neural bases of speech external feedback monitoring systems. Finally, the fact that the 4– 8-Hz brain → speech coupling was significantly enhanced during reading (compared to listening) also suggests that the brain does generate internal representations of self-produced syllabic sounds, as put forward by the predictive coding theory (Friston, 2010). Importantly, the motor origin of this effect supports the notion that, in this frequency band, the brain computes the time-course of the to-be-produced articulation.

### 4.3. Methodological considerations

First, there was no difference between *listen* and *playback* conditions in any of the tested aspects of speech brain tracking. This implies that the effects we uncovered (i) were not influenced by priming about upcoming speech content (intrinsic to *playback*) and (ii) not linked to a difference in speech rhythm between *listen* and *read*.

Second, neurophysiological mechanisms involved in overt language production are typically difficult to explore using MEG due to multiple sources of high-amplitude artifacts (e.g., head and jaw movements, muscular activity, etc.) that contaminate brain signals (see, e.g., Simmonds et al., 2014). Here, we used tSSS, ICA and threshold-based artifact rejection to remove these artifacts from brain signals. We then reconstructed brain activity with a minimum variance beamformer, an approach that specifically passes activity coming from locations of interest while cancelling external interferences (Hillebrand et al., 2005). Still, sensor and source speech brain tracking in the production condition indicated the presence of remaining movement artifacts characterized by coherence values comparable to genuine speech brain tracking/coherence values. It is therefore probable that these artifacts were mild and hence not suppressed by tSSS, ICA or beamforming.

Beyond attempting to suppress artifacts, we conducted two control analyses designed to evaluate the impact of remaining artifacts on our results. First, by computing the rPDC between speech signals and MEG signals at sensors with high amplitude artifacts, we could demonstrate that reading-induced artifacts spuriously inflate rPDC values in both directions. This supports our two main findings since reading (compared with listening) was associated with decreased <1 Hz tracking (rather than increased), and specifically increased 4–8 Hz tracking in the brain → speech direction (rather than in both directions). Finally, we used partial coherence analysis in sensor space wherein we subtracted the contribution of MEG signals at sensors on the edge of the sensor array to support our source-level results. This second control analysis corroborated the finding that 4–8 Hz tracking is enhanced during reading compared with listening. However, it suggested similar rather than lower level of <1-Hz tracking during reading compared with listening. Further studies based on artifact free electrophysiological signals (e.g., intracranial recording; Cogan et al., 2014) will be required to confirm source-space results. Also, we cannot exclude that the sources of 4–8 Hz tracking in the reading condition may have been shifted by the artifacts remaining in sensor data. Invasive electrophysiological recordings are warranted to identify the exact cortical network involved in tracking of self-produced speech, and specifically, to determine the relative contribution of STAC and parietal operculum.

Despite these limitations that warrant to take the results of this study with some caution, we demonstrate that the speech brain tracking observed at <1 Hz during *listen* and *read* is rather similar in terms of brain areas and tracking level. Furthermore, the results obtained at 4-8 Hz during *read* are in line with those previously reported by Ruspantini et al. (2012) during syllable production. These data therefore suggest the existence of common speech brain tracking phenomena during self-generated speech production accompanying reading aloud and perception while listening to somebody reading a text aloud. The generalization of these findings to production and perception of natural speech (e.g., during natural conversation) warrants further investigations. Still, this study represents a first step towards the understanding of the neural bases and functional aspects of speech brain tracking during speech production.

### 4.4. Conclusions

This study demonstrates that, during reading aloud, the reader’s brain tracks the slow temporal structure of the self-generated speech. The auditory cortex tracks phrases/sentences and the parietal operculum tracks syllables. Data also suggests that both tracking mainly engage feedback monitoring system, but with increased involvement of internal speech monitoring system for syllable tracking at different neocortical areas than those recruited during speech perception. In sum, this study brings unprecedented insights into how the human brain tracks the slow-temporal features of the auditory feedback during self-generation of speech.

## 5. Acknowledgments

Mathieu Bourguignon has been supported by the program Attract of Innoviris (grant 2015-BB2B-10), by the Spanish Ministry of Economy and Competitiveness (grant PSI2016-77175-P), and by the Marie Skłodowska-Curie Action of the European Commission (grant 743562). Nicola Molinaro has been supported by the Spanish Ministry of Economy and Competitiveness (grant PSI2015-65694-P), the Agencia Estatal de Investigación (AEI), the Fondo Europeo de Desarrollo Regional (FEDER) and by the Basque government (grant PI_2016_1_0014). Mikel Lizarazu has been supported by the Agence Nationale pour la Recherche (grants ANR-10-LABX-0087 IEC and ANR-10-IDEX-0001-02 PSL). Xavier De Tiège is Post-doctorate Clinical Master Specialist at the Fonds de la Recherche Scientifique (F.R.S.-FNRS, Brussels, Belgium).

This research was supported by the Basque Government through the BERC 2018-2021 program and by the Spanish State Research Agency through BCBL Severo Ochoa excellence accreditation SEV-2015-0490. The MEG project at the CUB Hôpital Erasme is financially supported by the Fonds Erasme.

